# Genomic signatures of island colonization in highly diverse primates

**DOI:** 10.1101/2024.12.13.628324

**Authors:** I. Colmonero-Costeira, K. Guschanski, M.L. Djaló, N. Fernandes, T. Camará, K.K.-H. Farh, L. Kuderna, J. Rogers, T. Marques-Bonet, M.W. Bruford, I-R.M. Russo, A. Jensen, M.J. Ferreira da Silva

## Abstract

Understanding how small populations cope with loss of genetic diversity and deleterious variation is crucial to address the current biodiversity crisis. Insular populations are particularly interesting as they have often persisted at low population sizes and higher inbreeding than their mainland counterparts. While the genome-wide consequences of inbreeding in threatened insular species have received some attention, comparative genomics between insular and mainland populations of wide-spread and genetically diverse species have rarely been performed. Yet, they are particularly well suited to inform about the consequences of drastic population declines from initially large populations - a phenomenon that is becoming increasingly common. The spot-nosed monkey (*Cercopithecus petaurista*), the Campbell’s monkey (*Cercopithecus campbelli*) and the green monkey (*Chlorocebus sabaeus*) are common and genetically diverse West African primates. Insular populations can be found at the Bijagós Archipelago, Guinea-Bissau. Here, we assessed the genome-wide diversity, inbreeding, genetic load and adaptive variation using whole genome sequencing data from insular and mainland populations. In the three species, island populations showed lower genome-wide diversity and higher inbreeding. Genetic drift has likely promoted the conversion of masked genetic load into realized load without increased purging of deleterious variation. Additionally, we found no evidence for accumulation of deleterious variation, suggesting that these populations are not yet at risk of extinction by genetic factors and may act as reservoirs of mainland genetic diversity. We highlight, however, that other anthropogenic factors are threatening these insular primates and therefore conservation management should target their immediate threats and safeguard against additional loss of diversity.

## Introduction

In a time when anthropogenic activities increasingly impact ecosystems, conservation of biodiversity has become a global challenge. Intraspecific genetic diversity has recently received wider recognition by governmental bodies (CBD, 2020; Hoban et al., 2021; Mastretta-Yanes et al. 2023). Since its inclusion in the Convention on Biological Diversity’s (CBD) Kunming-Montreal Global Biodiversity Framework, greater accessibility to genomic data for non-model species has improved our capacity to disentangle neutral and adaptive components of genetic diversity, opening the door to better understand species demographic and adaptive responses to environmental or anthropogenic changes (Allendorf et al., 2010).

Insular populations can be particularly vulnerable to extinction due to genetic factors resulting from the demographic processes inherent to the colonization dynamics. Island colonization typically involves a small number of founding individuals, which carry only a fraction of the total genetic diversity of the source populations (Allendorf et al., 2013; Martin et al., 2023). The intrinsic low levels of genetic diversity in insular populations tend to be aggravated by restricted or no gene flow with the source populations and increased inbreeding among founding individuals and their descendants. Genetic drift is stronger in populations with small effective population size (*N*_e_), as often is the case in insular systems (Allendorf et al., 2013). Although genetic drift is a neutral evolutionary process, it can also incur fitness costs by increasing the allele frequencies of loci with deleterious variation (Bertorelle et al., 2022; Dussex et al., 2023). As masked genetic load (i.e., recessive deleterious mutations present in their heterozygous state) is converted into realized load (i.e., homozygous deleterious mutations with expressed fitness effects; Bertorelle et al., 2022), individual genetic load may increase, reducing the individual fitness and increasing the risk of inbreeding depression (Bertorelle et al., 2022; Dussex et al., 2023; Smeds & Ellegren, 2023). However, the increased frequency of strongly deleterious mutations in their homozygous states makes them visible to selection, allowing for their removal through purging (Dussex et al., 2021; Grossen et al., 2020; Khan et al., 2021; van der Valk et al., 2024; von Seth et al., 2022; Xue et al., 2015). Although purifying selection is expected to purge mutations with large fitness effects, strong genetic drift can reduce the efficiency of selection in small populations, potentiating the gradual accumulation of mildly deleterious mutations during prolonged bottlenecks (Dussex et al., 2023). Such accumulation can generate a decline in fitness and exacerbate the extinction risk of small, isolated populations (Bataillon & Kirkpatrick, 2000; Khan et al., 2021).

Insular populations frequently exhibit genetic and phenotypic differences compared to mainland populations in response to novel ecosystems. This phenomenon, collectively known as the “island syndrome”, often includes changes in morphology, behavior and life-history traits (Adler & Levins, 1994). It has been attributed to adaptive processes in response to the new habitats where different traits, more advantageous to the insular environment, are selected for (Martin et al., 2021; Payseur & Jing, 2021; Sendell-Price et al., 2020; Welles & Dlugosch, 2019). Another hypothesis is that loci under strong selection on the mainland become effectively neutral on the islands due to the relaxation of natural selection (Cui et al., 2021; Wang et al., 2023). Consequently, non-synonymous variants can drift towards fixation in the insular populations.

Due to their vulnerability coupled with high potential for neutral and adaptive differentiation, insular populations are of great conservation concern. Previous studies have focused mainly on highly threatened insular populations from already genetically depauperate species (e.g., kākāpō, *Strigops habroptilus;* Dussex et al., 2021; Chatham Island black robin, *Petroica traversi*, von Seth et al., 2022). However, little attention has been given to widespread and genetically diverse species. Such species, many of which are classified as non-threatened on the International Union for Conservation of Nature (IUCN) Red List, are becoming increasingly threatened due to anthropogenic activities in their ranges (Finn et al., 2023). As the genomic consequences of inbreeding and decreasing population sizes of widespread populations may differ from those of more threatened taxa (Robinson et al., 2016; Taylor et al., 2024), it is increasingly important to consider non-flagship species in the field of conservation genomics in order to pre-emptively promote their conservation.

Here, we investigated the effects of insularity and low population size in three guenon species, tribe Cercopithecini, the most genetically diverse African primate group (Jensen et al., 2023; Kuderna et al., 2023). We used sequencing data to compare genome-wide diversity, inbreeding and genetic load between insular and mainland populations of the spot-nosed monkey (*Cercopithecus petaurista*, Scheber, 1774), the Campbell’s monkey (*Cercopithecus campbelli*, Waterhouse, 1838), and the green monkey (*Chlorocebus sabaeus*, Linnaeus, 1766) in Guinea-Bissau, West Africa. Although classified as non-threatened by the IUCN, our focal species are declining throughout their ranges (Gonedelé Bi et al., 2020; Matsuda Goodwin, et al., 2020; Matsuda Goodwin, et al., 2020). In Guinea-Bissau these are the most hunted primate species for bushmeat trade (Ferreira da Silva et al., 2021; Minhós et al., 2013) and the insular populations are not considered for conservation management by local authorities (Colmonero-Costeira et al., 2023).

## Methods

### Study area and focal species

The Bijagós Archipelago is a continental archipelago of 88 islands and islets located off the coast of Guinea-Bissau, West-Africa. It has been proposed that the separation of the archipelago from mainland started as early as ca. 19,000 years ago (YA) but is thought to have become fully isolated during the Holocene, particularly during the lower Flandrian transgression, ca. 6,000 – 12,000 YA (Alves, 2007). Both climate and vegetation are similar to southern areas of the mainland (Catarino et al., 2008), which is characterized as a mosaic of forest of Guinean affinity and woodland savannah. Primates coexist with local communities whose arrival on the archipelago has been associated with the expansion of West African trading empires, including the peak expansion of the Ghana and Mali Empires, c.a. 800 – 1,200 YA (Rodney, 1970 in Lundy, 2015).

The spot-nosed monkey and the Campbell’s monkey are classified by the IUCN as Near Threatened (Matsuda Goodwin, et al., 2020; Matsuda Goodwin, et al., 2020), whereas the green monkey is listed as Least Concern (Gonedelé Bi et al., 2020). These guenon species are the only non-human primates occurring on the Bijagós Archipelago (Colmonero-Costeira et al., 2019; Gippoliti & Dell’Omo, 2003). Across the archipelago, the distribution of the three species does not overlap except on Caravela Island, where both the spot-nosed monkey and the Campbell’s monkey can be found in sympatry (Colmonero-Costeira et al., 2019; Gippoliti & Dell’Omo, 2003). The three species are widespread and considered habitat generalists, exhibiting high behavioral and dietary flexibility (Bersacola et al., 2022; Rowe & Myers, 2016).

In Guinea-Bissau, the spot-nosed monkey (*santchu nariz-branku* or *santchu-bidjugu* in Creole) is possibly restricted to the Bijagós Archipelago (Gippoliti & Dell’Omo, 2003). Historically, the species is thought to have been present on the mainland (Gippoliti & Dell’Omo, 2003) but has been absent/not reported from the mainland for the last 30 years (Bersacola et al., 2018, 2022; Colmonero-Costeira, *unpublished*). The Campbell’s monkey (*kankulma* or *santchu-mona* in Creole) and the green monkey (*santchu di tarrafi* in Creole) are considered the most abundant non-human primate species in the country (Bersacola et al., 2018; Karibuhoye, 2004). However, past studies suggest that their populations are threatened by illegal commercial hunting and habitat destruction (Gippoliti & Dell’Omo, 2003). Campbell’s and green monkeys are the most traded primates for meat consumption in the urban bushmeat markets in the capital city and the most commonly available species in bushmeat dedicated restaurants/bars in the south of the country (Ferreira da Silva et al., 2021; Minhós et al., 2013). On the Bijagós Archipelago, the commercial trade of spot-nosed monkey meat has been recently described (Colmonero-Costeira et al., 2023) and subsistence hunting is reported for the remaining two species (see below).

### Sample collection

Tissue samples of spot-nosed monkeys, Campbell’s monkeys, and green monkeys were obtained as part of previous research that focused on primate meat hunting and consumption in Guinea-Bissau between 2010 and 2017. Samples from the Bijagós Archipelago were collected directly from local hunters or villagers preparing the carcasses for consumption (2002 – 2016). Samples from mainland Guinea-Bissau were collected at meat markets in the capital city, Bissau (March – June 2010; Minhós et al., 2013), or from drinking establishments located in the vicinity of protected areas in southern mainland (2015 – 2017; Ferreira da Silva et al., 2021). The DNA extractions of samples collected in Bissau is described in Minhós et al. (2013). For the other samples, we extracted the DNA of the samples collected between 2015 and 2017 using a salting out procedure (Miller et al., 1988) at the CIBIO-InBIO research centre. Negative controls were included throughout the entire process to test for DNA contamination. Samples with low DNA concentrations were re-extracted using the Qiagen DNeasy Blood & Tissue kit (QIAGEN® Hilden, Germany) following the manufacturer’s protocol. To minimize the inclusion of highly related individuals in the dataset, we selected samples collected in different weeks or different commercial establishments or in different hunting events. A total of 50 samples were sequenced on Illumina NovaSeq6000 (150bp PE), aiming for 30x coverage per sample. We generated high quality whole-genome sequences from the three guenon species present on the Bijagós Archipelago (eight spot-nosed, one Campbell’s and three green monkeys), and the mainland counterparts of Campbell’s (n = 25) and green monkey (n = 10). (Supplementary data S1). Since no spot-nosed monkeys are present on the mainland Guinea Bissau, we included a publicly available spot-nosed monkey genome of unknown geographic origin from mainland Africa in the analyses (Kuderna et al., 2023).

### Mapping and variant calling

We used Picard v2.23.4 (http://broadinstitute.github.io/picard/) to add read group information and mark adapter content in the sequencing reads and aligned them to the rhesus macaque (*Macaca mulatta*) reference genome (Mmul_10, GCF_003339765.1) with the Burrows-Wheeler Aligner (BWA) v0.7.17 (Li & Durbin, 2009). Picard was used to sort and deduplicate the mapped bam-files. Next, we used GATK v4.3.0.0 to call genotypes per sample with HaplotypeCaller, which were subsequently combined with CombineGVCFs and jointly genotyped with GenotypeGVCFs. We excluded insertions and deletions, and used VariantFiltration to filter variants based on the following exclusion criteria: QD < 2.0, QUAL < 30.0, SOR > 3.0, FS > 60.0, MQ < 40.0, MQRankSum < -12.5, ReadPosRankSum < -8.0. Repetitive regions were excluded following the SNPable regions pipeline (https://lh3lh3.users.sourceforge.net/snpable.shtml). We also masked heterozygous genotypes with skewed allelic balance (minor allele support < 0.25), as well as genotypes with less than half or more than double the genome-wide average read depth per sample.

### Mitochondrial genome assembly and haplotype networks

The mitochondrial genomes were assembled using the MitoFinder pipeline (Allio et al., 2020). We started the assembly by trimming adapter sequences using Trimmomatic (Bolger et al., 2014) and running MitoFinder with the metaspades assembler. The assembled contigs were corrected for circularization and rotated to the same starting position using a custom Python script (https://github.com/axeljen/guenon_phylogenomics) and annotated using the green monkey (NC_008066.1) reference mitochondrial genome. Annotated assemblies were aligned for each species separately using MAFFT aligner (Katoh et al., 2002). We inspected the species-specific alignments visually and trimmed the non-coding hypervariable region 1 (positions 16,010 – 16,361), and hypervariable region 2 (positions 56 – 364) of the green monkey reference genome and misassembles/misalignments. For the spot-nosed monkey we included publicly available mitochondrial genomes (JQ256931.1, JQ256932.1, JQ256982.1 and JQ256983.1) in the final alignment (Guschanski et al., 2013). We visualized the relationships between the assembled mitochondrial genomes by reconstructing median joining haplotype networks using PopART (Leigh & Bryant, 2015).

### Population structure and historical population size

To assess population structure between the insular and mainland populations, we conducted a principal component analysis (PCA) in PLINK v1.9 (Purcell et al., 2007), removing positions with missing genotypes within each species (--geno 0) and thinning the total autosomal SNPs to 100,000 SNPs. We inferred the long-term changes in *N*_e_ of all the populations using the pairwise sequentially Markovian coalescent model implemented in beta-PSMC (Liu et al., 2022) with default parameters and five sub-atomic time intervals. The variance of the estimated *N*_e_ trajectories was assessed by 20 randomized resampling runs across the genome. We scaled the output assuming a generation time of 11 years for the spot-nosed monkey, 12 years for both Campbell’s and green monkeys, and a mutation rate of 4.65x10^-9^ per site per generation (average mutation rate across guenons; Kuderna et al., 2023).

### Genome-wide diversity and inbreeding

We identified autosomal runs of homozygosity (ROH) using PLINK v1.9 and estimated autosomal heterozygosity for each individual genome using vcftools (Danecek et al., 2011). To minimize the detection of false positive ROH, we scaled the number of SNPs that constituted a ROH (--*homozyg-snp* setting) by the within species average heterozygosity (Purfield et al., 2012):

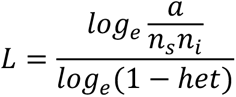

where n_s_ is the number of genotyped SNPs, n_i_ the number of genotyped individuals, α the false positive ROH threshold (set at 0.05), and *het* the mean heterozygosity. Based on L, the minimum number of SNPs that constituted a ROH was set to 77, 88, and 80 for spot-nosed monkey, the Campbell’s monkey, and the green monkey, respectively. We selected a scanning window length equal to L as suggested by (Meyermans et al., 2020). For the remaining ROH search parameters we used the default values. The fraction of the genome contained in ROH (F_ROH_) for each individual was estimated as:

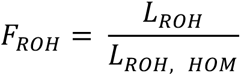

where *L_ROH_* is the sum of the length of all detected ROHs, and *L_ROH, HOM_* is the maximal detectable ROH length of a synthetic homozygous genome of each individual, under the same ROH searching parameters (Meyermans et al., 2020). The synthetic homozygous genome for each of the individuals was obtained by changing all heterozygous genotypes to homozygous using a custom Python script. This minimized the impact of stochastic genome-specific sequencing artefacts that potentially prevent the correct detection of ROH in window-based methods (Meyermans et al., 2020).

We estimated the distribution of ROHs arising from inbreeding after the isolation of the Bjagós Archipelago or during human coexistence using the expected ROH coalescent time in generations (*g*):

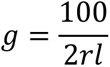

equivalent to:

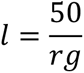

where *r* is recombination rate and *l* the ROH length (Thompson, 2013). We used the rhesus macaque (*Macaca mulatta*) average genome-wide recombination rate of 0.448 cM/Mb (Xue et al., 2020). As such, we established ROH ≥ 1 Mb as human-contemporary (up to 110 guenon generations, ≈ 1,200 YA) and ROH <1 Mb as historical (older than 110 guenon generations ago). The chosen time windows reflect the earliest time of the arrival of humans to the Bijagós Archipelago and the isolation of the archipelago, respectively.

### Genetic load

To investigate if isolation on the islands may have led to the purging of deleterious mutations, we characterized the derived positions in coding regions across the insular and mainland guenon genomes using Ensembl’s variant effect predictor (VEP; McLaren et al., 2016). Prior to the VEP annotation, we polarized the ancestral alleles using the Guinea baboon (*Papio papio*; GCA_028645565.1), the Angolan colobus monkey (*Colobus angolensis*; GCF_000951035.1), and rhesus macaque (Mmul_10) as outgroups. The additional outgroups were aligned to the Mmul_10 reference genome using Minimap2’s cross-species full-genome alignment (Li, 2018), assuming a species divergence below 20% (asm20). We called the genotypes on the outgroups using bcftools mpileup, multiallelic-caller (-m) (Danecek et al., 2021). Alleles present in the three outgroup species in the homozygous state were defined as ancestral. These positions were identified using a custom Python script. Invariable sites in the guenons were removed for downstream analysis.

The effects of the retained variants were predicted based on the *Macaca mulatta* (Mmul_10) annotations. For sites on overlapping transcripts, we selected a single prediction per variant based on VEP’s standard ordered set of criteria (--pick), including the canonical status of a transcript, APPRIS functional isoform notation (most functionally important transcripts), transcript support level, biotype (protein coding preferred), consequence rank (most severe). We classified the annotated variants as loss-of-function (LoF: transcript ablation, stop gained, stop lost, start lost, frameshift variant, splice region variant, splice donor variant, splice acceptor variant), missense, and synonymous. Missense variants were further classified into two classes: 1. likely deleterious or 2. tolerated, using the Experimental exchangeability score of amino acids (EX; Yampolsky & Stoltzfus, 2005). We calculated the EX of missense variants using a custom Python script adapted from Smeds and Ellegren (2022) (https://github.com/linneas/wolf-deleterious). Sites were classified as likely deleterious if the Experimental exchangeability score was ≤ 0.256 (Williamson et al., 2005).

A decrease in the number of deleterious variants across the genome can potentially arise as a combination of purifying selection and stochastic loss due to genetic drift. Considering that without selection, genetic drift does not change allele counts of the derived mutations (Dussex et al., 2023), we estimated the individual genetic load as the relative abundance (RA) of the derived alleles (*A^X^_A_*) for each individual (*i*) of population *x* and standardized it by the number of derived alleles within species across the different mutational categories:

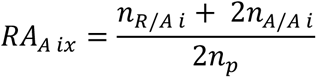

where *n_R/A i_* is the number of derived heterozygous positions (R, reference allele; A, alternative allele), *n_A/A I_* is the number of derived homozygous positions and *n_p_* is the total number of derived positions across all the genomes for a particular species (insular and mainland).

Furthermore, based on the VEP output, we performed a functional analysis of the LoF variants that have become fixed in the homozygous state in insular populations. We conducted a gene ontology (GO) enrichment analysis using g:Profiler (https://biit.cs.ut.ee/gprofiler/gost) and the g:SCS multiple testing correction algorithm on these sets of genes (Kolberg et al., 2023).

### Differential variation in protein-coding genes

We identified variants in protein-coding regions that were fixed for alternative alleles between insular and mainland populations of the same species using a custom Python script (Jensen et al., 2023). We annotated the differentially fixed positions using VEP. To filter out variants that accumulate through drift in potential pseudogenes, we removed all genes that contained a LoF variant. Next, we looked for sets of genes that contained differentially fixed variants in at least two of the studied guenon species and conducted a GO enrichment analysis using g:Profiler.

## Results

### Sequencing and genotyping

Across the 48 genomes, average autosomal mapping coverage of the dataset varied between 25.95x and 35.00x (median = 30.35x; Supplementary data S1). After calling the genotypes against the rhesus macaque Mmul_10 reference genome, we identified 18,415,827 autosomal biallelic SNPs in the spot-nosed monkey, 22,007,478 SNPs in the Campbell’s monkey, and 15,760,456 SNPs in the green monkey. Additionally, we assembled and annotated the mitochondrial genomes for all the samples.

### The three insular populations are differentiated from mainland populations and show reduced long-term *N*_e_

The PCAs suggested the existence of population structure in all three guenon species, with PC1 differentiating insular and mainland populations within each species and explaining 33.3%, 7.19% and 15.7% of the variance for the spot-nosed, Campbell’s, and green monkeys, respectively (Fig. 1). Further differentiation was observed between the spot-nosed monkey individuals from Canhabaque and Caravela Islands. In contrast, the green monkeys from Ganogo and Orango Islands (separated by a ∼500 m water channel) were almost indistinguishable from each other in the PC space. In the absence of clear population structure, individuals from these two neighboring islands were treated as one population in downstream analyses, designated as the Orango Islands Group (OIG). Mainland samples were generally more spread along the PC2 axis indicating more variation.

**Fig. 1.**
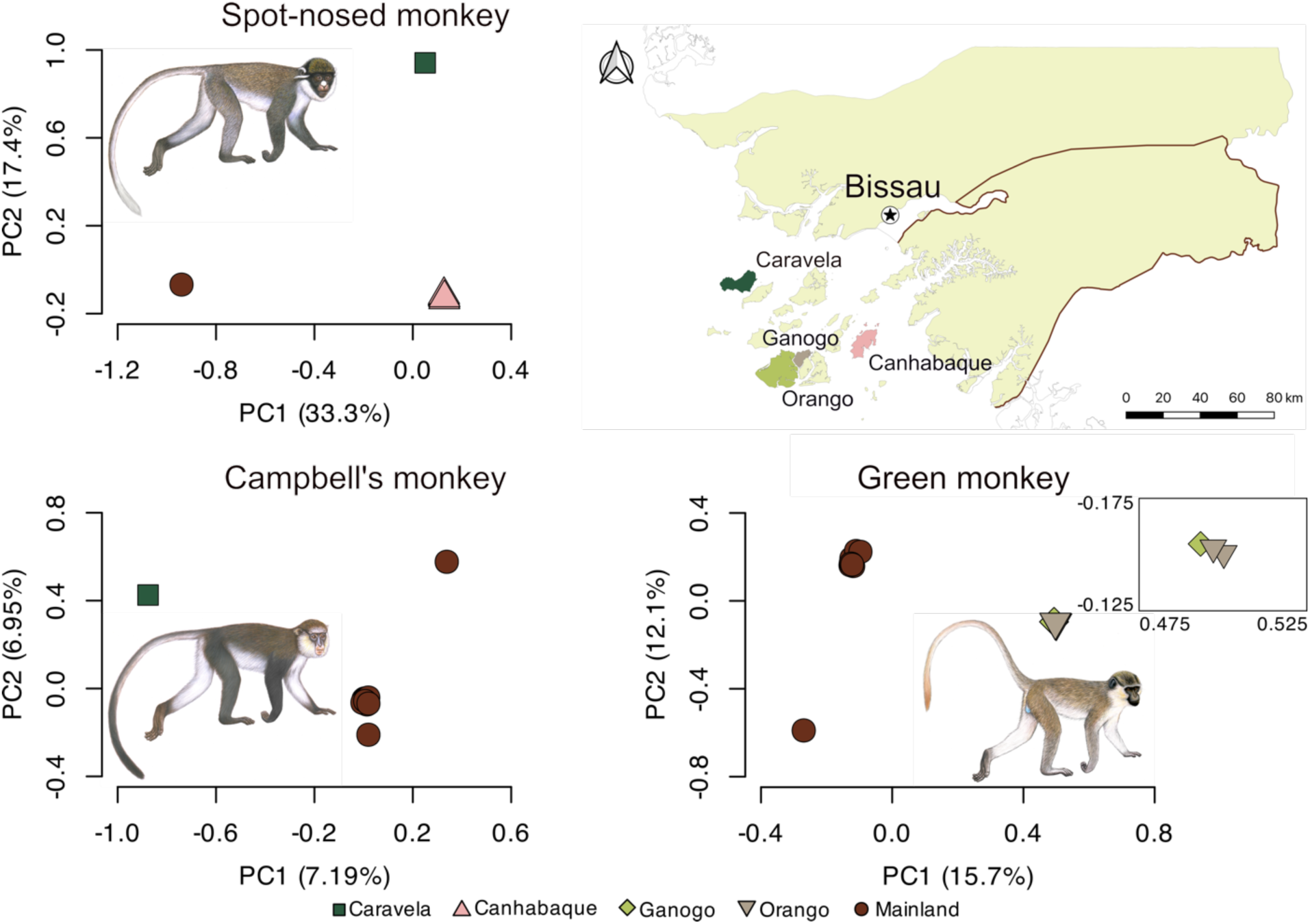
Principal Component Analysis (PCA) based on autosomal biallelic single nucleotide polymorphisms. An inset focusing on the insular green monkeys is shown on the right of the green monkey PCA plot. Each individual sample is colored according to their sampling area, as illustrated on the map of Guinea-Bissau, West Africa. The exact provenance of the mainland samples of Campbell’s and green monkeys is not known but they likely originate from Southern Guinea-Bissau (represented by the brown contour in the map in the upper right side; Minhós et al., 2013; Ferreira da Silva et al 2021). The geographic origin of the mainland spot-nosed sample is unknown, but it likely belongs to the eastern subspecies based on mtDNA analyses (Fig. S2). Illustrations copyright 2022 Stephen D. Nash / IUCN SSC Primate Specialist Group. Used with permission.

None of the mtDNA haplotypes sampled from the islands were shared with the mainland in any of the three species (Fig. S1 – 4). In the Campbell’s monkey and the green monkey, the pairwise number of substitutions between insular and mainland haplotypes were within the range of the pairwise comparisons among the mainland haplotypes. In contrast, the spot-nosed monkey showed insular haplotypes that differed from each other by only a few substitutions, but these haplotypes were separated from the mainland haplotype by more than 400 substitutions. Since the geographic origin of the mainland spot-nosed individual is not known, this separation could be the result of deep population structure within the species, such as genetic differentiation at the sub-species level. After the inclusion of the historical mtDNA genomes (Guschanski et al., 2013), the mtDNA haplotype from the mainland spot-nosed monkey included in the dataset was more closely related to mtDNA haplotypes sampled in Togo, Liberia and Ivory Coast. This result suggests the mainland individual used in this study could be from the eastern *C. petaurista* subspecies (*Cercopithecus petaurista petaurista*, Jentink, 1886), whereas the archipelago individuals likely belong to the western subspecies (*C. p. buettikoferi*), which would explain the deep genomic divergence.

We estimated changes in *N_e_* over time using beta-PSMC analyses. Overall, the three species showed different demographic histories roughly before the onset of the Penultimate Glacial Period (PGP, > 200,000 YA; Fig. 2). The maximum *N_e_* of the three species was observed close to the end of the PGP (slight timing variation between species), after which they all underwent a severe decline in *N*_e_ that extends until the end of the Last Glacial Period (LGP, ∼12,000 YA). The insular populations showed more rapid and severe declines than the mainland populations since the PGP, with population trajectories diverging between 100,000 – 40,000 YA in the three species.

**Fig. 2.**
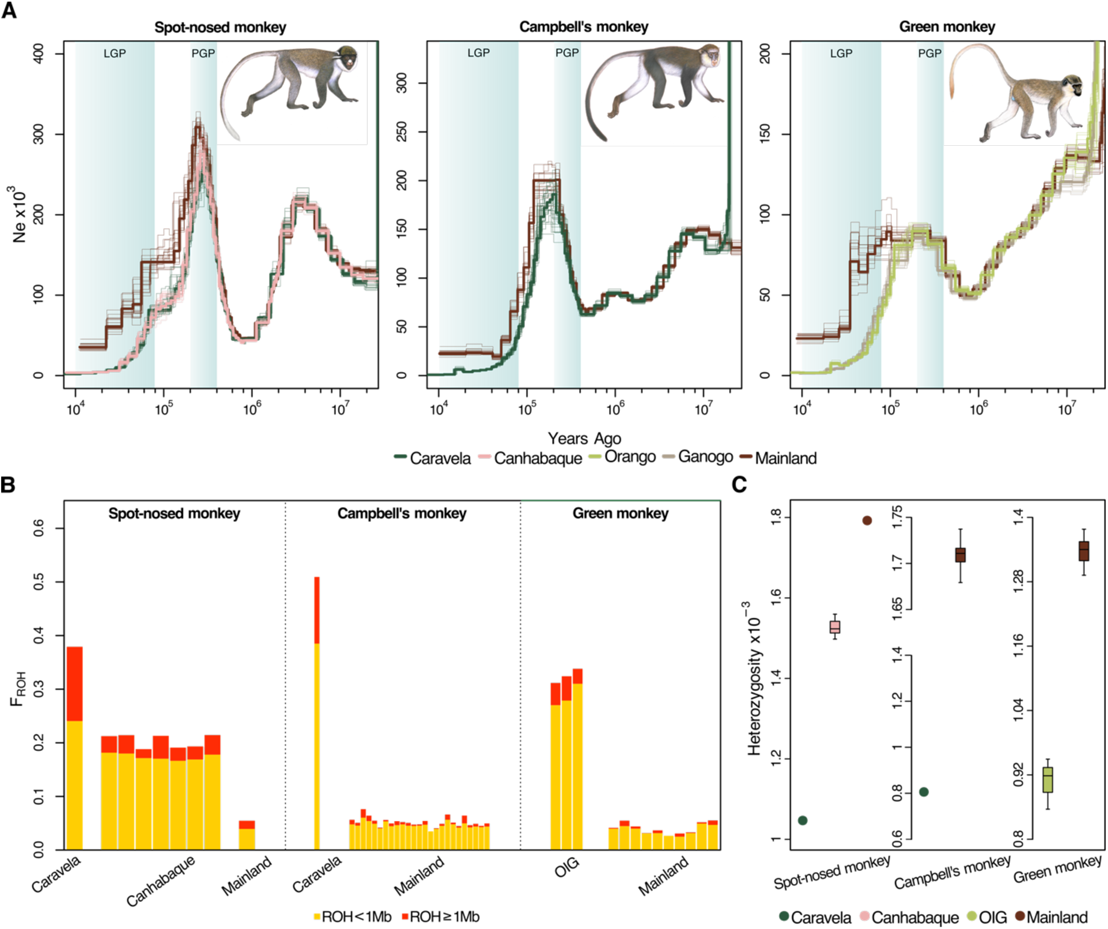
Genome-wide consequences of insularity in spot-nosed (left), Campbell’s (center), and green monkey populations (right). A) Estimation of long-term changes in *Ne* of populations using the pairwise sequentially Markovian coalescent model of insular (Caravela, Canhabaque, Orango and Ganogo) and mainland populations as inferred using beta-PSMC (Liu et al., 2022). Colors correspond to the sampling locations. Bootstrap runs are represented as pale lines. Light blue backgrounds correspond to the Last Glacial Period (LGP) and the Penultimate Glacial Period (PGP). For illustrative purposes, only a single genome per population is represented as the variation within populations is negligible (Fig. S5). B) Faction of the genome in ROH (FROH). FROH for each sampled individual is represented in the stacked bars. The proportion of short ROHs (< 1 Mb, 111 – 1,450 guenon generations, ≈ 1,211 – 16,000 YA) and long ROHs (≥ 1 Mb, 110 generations ago, ≈ 1,200) are represented in yellow and red, respectively. C) Heterozygosity based on bi-allelic autosomal SNP positions. The circles or boxplots (when N > 3 genomes) were colored according to the sampling location. Differences in autosomal heterozygosity between insular and mainland green monkeys were significant [(Welch’s two sample t-test, p<0.001; statistical significance could not be tested for the other species (N < 3 genomes in one of the groups)]. Green monkey individuals from Ganogo and Orango islands were clustered together into the Orango Islands Group (OIG). Note differences on the y-axis for each species. Illustrations copyright 2022 Stephen D. Nash / IUCN SSC Primate Specialist Group. Used with permission.

### Insular populations show signatures of elevated inbreeding and loss of genetic diversity

The insular individuals of the three species showed ∼73 – 90% higher proportion of the genome in ROH than mainland populations (Fig. 2B). The differences between insular and mainland populations were driven by both an increase in short (< 1Mb) and long ROHs (≥ 1Mb). The largest differences in the proportion of the genome in long ROHs was found in the spot-nosed monkey and Campbell’s monkey from Caravela Island, which displayed a ∼90% and 98% increase in ROH compared to the mainland individuals of the same species, respectively. Moreover, all insular individuals displayed lower heterozygosity (∼15 – 53% fewer autosomal heterozygous positions, Fig. 2C).

### Island populations do not show increased purging nor accumulation of deleterious variants

To understand if small effective population sizes and increased inbreeding associated with insularity led to changes in the frequency of deleterious mutations, we estimated the individual genetic load for each genome. After identifying the ancestral states of the segregating SNPs in the guenon species using three outgroup species, a total of 181,372 derived SNPs in the spot-nosed monkey, 217,297 in the Campbell’s monkeys, and 189,118 in green monkeys, were annotated (Table S1). In general, insular and mainland populations showed similar patterns across all categories of derived variants. In all three species, we found that the proportion of homozygous derived variants is higher in island individuals than in mainland individuals regardless of variant category (Fig. 3). This is a pattern consistent with conversion of masked into realized genetic load in the insular populations by inbreeding and genetic drift. There was a general trend of lower numbers of LoF and other deleterious variants among insular individuals compared to their mainland conspecifics (Table S1). However, after standardizing for the relative abundance of segregating alleles within species, there were no apparent differences between islands and mainland individuals (Fig. 3). Additionally, we highlight that the patterns between the genetic load of insular and mainland individuals across different categories of deleterious variants were similar to those of synonymous variants, which are expected to behave neutrally (Fig. 3).

**Fig. 3.**
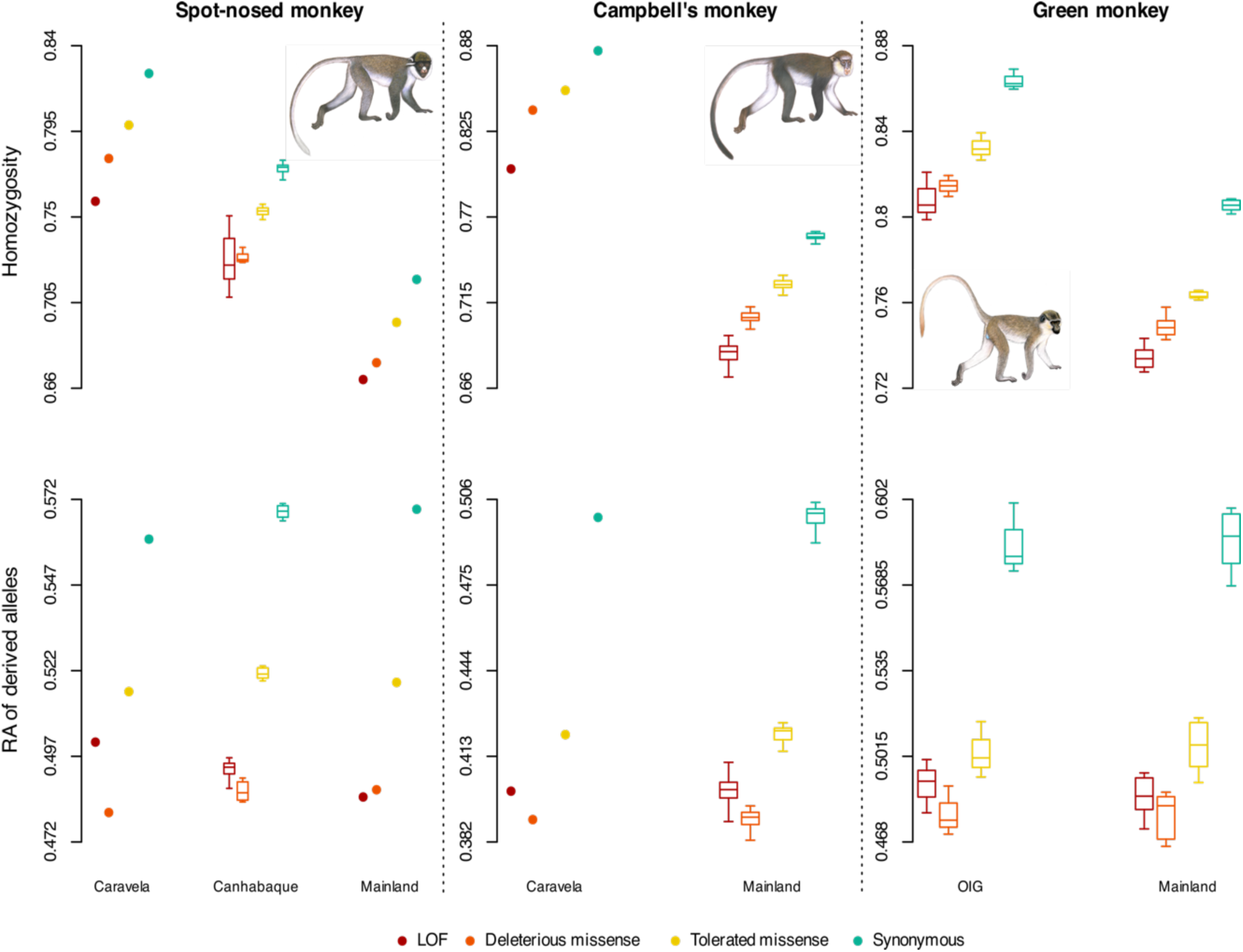
Individual mutational load in insular spot-nosed, Campbell’s, and green monkey populations. Upper panels: Homozygosity of variants across different categories of mutational effects of the annotated derived SNPs. Lower panels: Relative abundance of derived alleles. Boxplots are presented when N>3 individuals. The statistical comparisons for both the homozygosity and the relative abundance of derived alleles was performed within each variant category (i.e., LoF insular vs. LoF mainland) for the green monkey [Welch’s two sample t-test: homozygosity p < 0.01 across all categories; relative abundance of derived alleles, 0.32 < p < 0.90 across all categories; statistical significance could not be tested for the other species (N < 3 genomes in one of the groups)]. Missense mutations were classified as potentially deleterious or tolerated based on the experimental exchangeability score (Yampolsky et al., 2005).

In the insular spot-nosed monkeys we identified a total of 17 genes containing LoF variants that have become fixed in homozygous states, 32 genes in the insular Campbell’s monkey, and eight genes in the insular green monkeys (Supplementary data S2). These sets of genes were species-specific, with some being associated with immune response (spot-nosed monkey: APOL5; Campbell’s monkey: ACP5, ADGRG5, TRBV7-3, REG3G; green monkey: IFI16) and fecundity (Campbell’s monkey: PADI6). Gene ontology (GO) enrichment analysis revealed no overrepresented GO terms.

### Differential variation in protein-coding genes between island and mainland populations is enriched for developmental processes

To explore the hypothesis that insularity can lead to specific, potentially adaptive, genetic and phenotypic changes between insular populations and their mainland counterparts, we identified all differentially fixed positions between the island and mainland populations in each species. We found 2,794 differentially fixed variants in protein-coding genes for the spot-nosed monkey, 491 for the Campbell’s monkey, and 125 for the green monkey. Within these gene sets, 203 genes were found in at least two species (Fig. S6). The GO enrichment analysis of this overlapped gene set revealed several overrepresented terms, mostly related to various metabolic functions and organism developmental processes (Supplementary data S3). Specifically, several of the genes with differential variation between insular and mainland populations are involved in cellular division and organogenesis (PDGFRA, TGFB2, E2F8, NIN), neuronal and muscular development (SYNE2, BRINP1, TNC, ARNT2, TENM4, ACTN3, FMN1), skeletal development (CHST11, COL2A1) and, thyroid development and functioning (PAX8).

## Discussion

Insular populations are often of conservation concern as they are typically more susceptible to loss of genetic diversity due to demographic processes associated with island colonization. In this work we explored the genomic signatures of insularity and reduced population size in guenons, a primate group known for their overall high genetic diversity (Jensen et al., 2023; Kuderna et al., 2023). By comparing island and mainland populations in three guenon species from the Bijagos Archipelago, Guinea-Bissau (in West Africa), we explored their common genetic patterns which likely represent the broad effects of insularity. Finally, we use our results to inform future conservation management actions for the three guenon species of the Bijagós Archipelago in Guinea-Bissau. Our results are potentially applicable to other wide-spread and genetically diverse taxa, now under increasing threats from anthropogenic activities.

### Genome-wide effects of insularity

In all three guenon species considered here, island populations were genetically differentiated from the mainland populations. The timing of the divergence between insular and mainland populations varied between species but has generally happened between the PGP and the start of the LGP (250,000 – 100,000 YA), according to bPSMC analyses (Fig. 2A). Since this time period, the island populations have experienced a steadier decline in the estimated *N_e_* than their mainland counterparts. The observed lower genome-wide heterozygosity and higher F_ROH_ in the insular populations of all three species compared to mainland populations support long-term reduction of the *N*_e_ as suggested by the bPSMC. The genome-wide effects of population bottlenecks associated with island colonization could be further exacerbated in archipelagos, where sequential colonization events from the initial founding populations are likely (Allendorf et al., 2013; Martin et al., 2023). In our study species, this effect can potentially be observed in the spot-nosed monkey individual from Caravela, the outermost island of the Bijagós Archipelago. This individual has lower genome-wide diversity and higher F_ROH_ compared to the Canhabaque population.

The timing of divergence between the insular and mainland populations as estimated by the bPSMC analyses is unexpectedly old considering the proposed timing of isolation of the Bijagós Archipelago during the Holocene (ca 12,000 – 6,000 YA; Alves et al., 2007). Previous works have shown that fluctuations in stair-case plots from coalescent-based methods can reflect true changes in *N*_e_, cessation of panmixia, or both (Mazet et al., 2016). Indeed, the timing of the divergence of the *N*_e_ trajectories coincides with cycles of dryer and wetter climate and, consequently repeated retraction and expansion of forests across sub-Saharan Africa (Hoag & Svenning, 2017) – a known promoter of changes in connectivity in forest dwelling species such as primates (van der Valk et al., 2024). Thus, it is plausible that the reported divergence of the *N*_e_ trajectories reflects a scenario of the disruption of panmixia/population structure arising prior to the colonization of the archipelago. Moreover, the mainland populations included here may not represent the most closely related source populations, particularly for the spot-nosed monkey, whose presence has not been reported from mainland Guinea-Bissau for over 30 years.

A recurring question in conservation genomics regarding the fate of small and inbred populations, such as those inhabiting insular systems, is whether genetic load is accumulating after a demographic bottleneck or if genetic purging is effective in decreasing the genetic load of highly deleterious mutations, thus preventing mutational meltdown. The decrease in the frequency of LoF variants following the conversion of masked load (i.e., heterozygous, partially recessive deleterious mutations) into realized load (i.e., homozygous states) is often the main diagnostic features of purging in bottlenecked populations (Dussex et al., 2021; Grossen et al., 2020; Khan et al., 2021; van der Valk et al., 2024; von Seth et al., 2022; Xue et al., 2015). Although a higher proportion of homozygous LoF variants was found in the insular guenon populations studied here, similar patterns were present in missense and neutral variants. Furthermore, after standardizing the genetic load to the number of segregating sites (i.e., relative abundance of derived alleles), we observed that insular and mainland individuals have similar numbers of deleterious alleles across all variant categories, which is expected under genetic drift alone (Dussex et al., 2023; Smeds & Ellegren, 2023).

Another commonly reported genomic consequence of prolonged periods of small *N_e_* is the accumulation of mildly deleterious (e.g., missense) mutations (Dussex et al., 2023). Since drift diminishes the effectiveness of purifying selection, *de novo* mildly deleterious mutations are more likely to drift to higher frequencies in small populations (Dussex et al., 2023). We do not observe such a pattern in the insular guenon populations, suggesting that the effectiveness of purifying selection against deleterious derived alleles has been retained overall (Dussex et al., 2023). We argue that the insular guenon populations of the Bijagós Archipelago lack the typical genomic features of small, inbred populations. Our results suggest that the increased inbreeding and loss of genetic diversity in the insular populations have, so far, not led to increased purging of highly deleterious variants nor accumulation of genetic load (Dussex et al., 2023; Robinson et al., 2016; Taylor et al., 2024).

Several guenon species have been successful in establishing insular populations following human-assisted introduction (e.g., green monkeys in Cape Verde (Almeida et al., 2024; Hazevoet & Masseti, 2011) and multiple Caribbean Islands (Denham & Denham, 1981), mona monkeys, *Cercopithecus mona,* in São Tomé and Príncipe and Grenada (Glenn & Bensen, 2013; Horsburgh et al., 2003)). Based on historical records, these populations are thought to have been founded by relatively small numbers of individuals and the populations have increased to thousands of individuals within 300 – 500 years. While most of these populations remain poorly studied, there are no reports suggesting the presence of deleterious inbreeding effects on the islands when compared to mainland Africa (e.g., Grenada mona monkeys; Glenn & Bensen, 2013). The success of guenon colonization and potential avoidance of inbreeding effects could be attributed to a high ecological and behavioral flexibility, and the relatively beneficial island ecosystems (e.g., decreased interspecific competition; Glenn & Bensen, 2013) and overall high genetic diversity (Jensen et al., 2023; Kuderna et al., 2023).

Finally, we observed differentially fixed variation between insular and mainland populations. Some of the affected genes were present in two or three species and encoded for metabolic and developmental processes, which included cellular proliferation and organ size. While the specific phenotypic expression of these genes is unknown, these functional categories are amongst those commonly associated with the “island syndrome” in vertebrate species (Nolte et al., 2020; Payseur & Jing, 2021; Sendell-Price et al., 202). As populations are exposed to novel ecological challenges in insular ecosystems, which include changes in inter-specific competition and predation dynamics, and access to novel food items, selection pressures acting in insular and source populations are likely to differ in several aspects (Welles & Dlugosch, 201). Common adaptations to the same insular ecosystem are a possible mechanism behind the overlapping gene sets across the three species (Payseur & Jing, 2021; Welles & Dlugosch, 201). However, since our results suggest that genetic drift is likely the predominant evolutionary mechanism differentiating insular and mainland populations, an alternative explanation is that these genes are under relaxed selection on the islands. The relaxation of selection in genes linked to increased fitness in the mainland populations would allow drift to fix non-synonymous variants that may be otherwise deleterious on the mainland (Cui et al., 2021; Wang et al., 202). We stress however, that these results should be interpreted with caution as differences in phenotypic traits between the guenon populations of the Bijagós Archipelago and the mainland populations have never been studied, the current mapping to a distant reference genome (rhesus macaque) and unbalanced sample sizes prevents the unbiased assessment of differentially fixed non-synonymous variation, as neutrally evolving variation could be wrongly annotated as protein coding variation (e.g., pseudogenes).

### Importance of insular populations for the conservation of genetically diverse species

Even though guenon populations of the Bijagós Archipelago are thought to be more threatened than their mainland counterparts - due to the overall loss of genetic diversity and inbreeding associated with insularity - this study did not find increased accumulation of deleterious variation on the islands. However, the increased inbreeding and genetic drift on the islands resulted in higher levels of realized load which may affect population fitness negatively. We found fixed LoF variants in genes involved in immune and reproductive functions in the insular populations of the three guenon species, although their fitness effects are unknown.

It is important to highlight that natural processes are not the only factor determining individual survival of these primates. Habitat degradation and harvesting of individuals for meat consumption are current conservation threats, which may increase their risk of extinction (Colmonero-Costeira et al., 2023). Indeed, the insular populations of the three species show significantly higher fraction of their genome in long runs of homozygosity, which likely arose due to mating of closely related individuals in the last 1,200 years after the arrival of humans on the archipelago. In particular, the illegal harvesting for the commercial meat trade has been described recently and is thought to be an increasing practice on the islands (Colmonero-Costeira et al., 2023). In the absence of immediate threats from genetic factors, long-term conservation of these insular populations should be promoted mainly by reducing their immediate threats and safeguarding from additional loss of diversity by anthropogenic factors. For Guinea-Bissau in particular, the insular guenon populations are of high conservation interest, particularly the spot-nosed monkeys. We show that these relict populations are not genetically depauperate and could potentially act as reservoirs for the recently extinct (or very rare) mainland populations, creating the opportunity for future re-introduction of the species in mainland Guinea-Bissau.

Recent studies have proposed that small and threatened populations that underwent extensive purging are expected to be less sensitive to the effects of additional bottlenecks and can tolerate smaller *N_e_* and still avoid severe inbreeding depression and extinction due to genetic factors (Caballero et al., 201). However, not much is known about the response of genetically diverse species to demographic bottlenecks, such as the insular guenons studied here. Akin to this system, high realized genetic load in the absence of purging following population bottlenecks has been reported in other widespread and diverse species, such as the North American caribou (*Rangifer tarandus*) (Taylor et al., 2022) and the Channel Island fox (*Urocyon littoralis*) (Robinson et al., 2016). In these populations the interplay between the increasing anthropogenic threats and the dynamics of deleterious variation following demographic declines remains poorly studied and ought to be further investigated in order to pre-emptively promote their conservation.

## Author contributions

Conceptualization: I.C-C., A.J., K.G., I.M.R., M.W.B., M.J.F.S.; Methodology and analyses: I.C-C., A.J.; Sample acquisition: M.L.D., N.F., I.C., M.J.F.S.; Funding acquisition: M.J.F.S., K-H.K.F., L.K., J.R., T.M-B., Writing—original draft: I.C-C; Writing—review and editing: all authors. A.J. and M.J.F.S. contributed equality to this work.

## Acknowledgments

We dedicate this work to Professor M.W. Bruford – our mentor, colleague and friend. His enthusiasm, guidance, and support were key to the success of this study and to the advancement of the field of conservation genetics of the primates in Guinea-Bissau. We would like to acknowledge the Guinea-Bissau governmental agency Instituto de Biodiversidade e Áreas Protegidas (IBAP), namely to the former director Dr A. Silva and Dr J. Biai, and to current director Dr A. Regalla, and the directors of protected areas and staff members – Dr Said A., Dr A. Cá, Dr. J. Mané and Dr S. Danfa, for permits and logistical support during fieldwork. We thank I. Espinosa and H. Foito for logistical support in Bissau. We are grateful to the local communities of the Bijagós Archipelago and from southern mainland Guinea-Bissau that facilitated the execution of this work. We thank Prof L. Palma for facilitating tissue samples of spot-nosed monkeys from the Bijagós Archipelago and logistical support. We acknowledge Dr R. Godinho and the CTM (Centre for Molecular Analysis, CIBIO-InBIO) staff for all the help and support with the molecular analysis of the tissue samples collected in 2015-2017.

## Ethics disclosure and benefits-sharing

All collected tissue samples used in this work were provided voluntarily and free of charge by local hunters and/traders. Informed consent to participate was obtained before beginning research. The providers of samples remained anonymous to local agencies and law enforcers. All sampling was carried out with the approval of the National Institute for Biodiversity and Protected Areas (IBAP). Direção Geral das Florestas e Fauna (DGFF) authorized transportation and issued CITES export permits for tissue samples. Instituto da Conservação da Natureza e Florestas (ICNF, Portugal) issued the CITES importation permits to Portugal (*Cercopithecus petaurista* – N.°18PTLX00592I and 23PTLX00512I, *Cercopithecus campbelli –* N.°18PTLX00590I, *Chlorocebus sabaeus* – N.°/No 18PTLX00586I). The Nagoya protocol was not in place in Guinea-Bissau at the time of sampling. However, we acknowledged the work of local researchers by including them in research decisions and disseminations of results and as co-authors of this manuscript. Results have been shared with the local governmental agencies through regular meetings and progress reports. Results of this work will be disseminated to the local communities through meetings and communications in local media. The authors have no competing interests to declare that are relevant to the content of this article.

## Data availability

The sequencing data used in this project are available on the European nucleotide archive (https://www.ebi.ac.uk/ena), under accession numbers XXX (provided upon acceptance). Custom scripts used for data analyses are available at https://github.com/Colmonero-CI/Guenons_POPGENOM.

