## Supplementary material for "Genomic signatures of island colonization in highly diverse primates": Fig. S1 - S6; Table S1

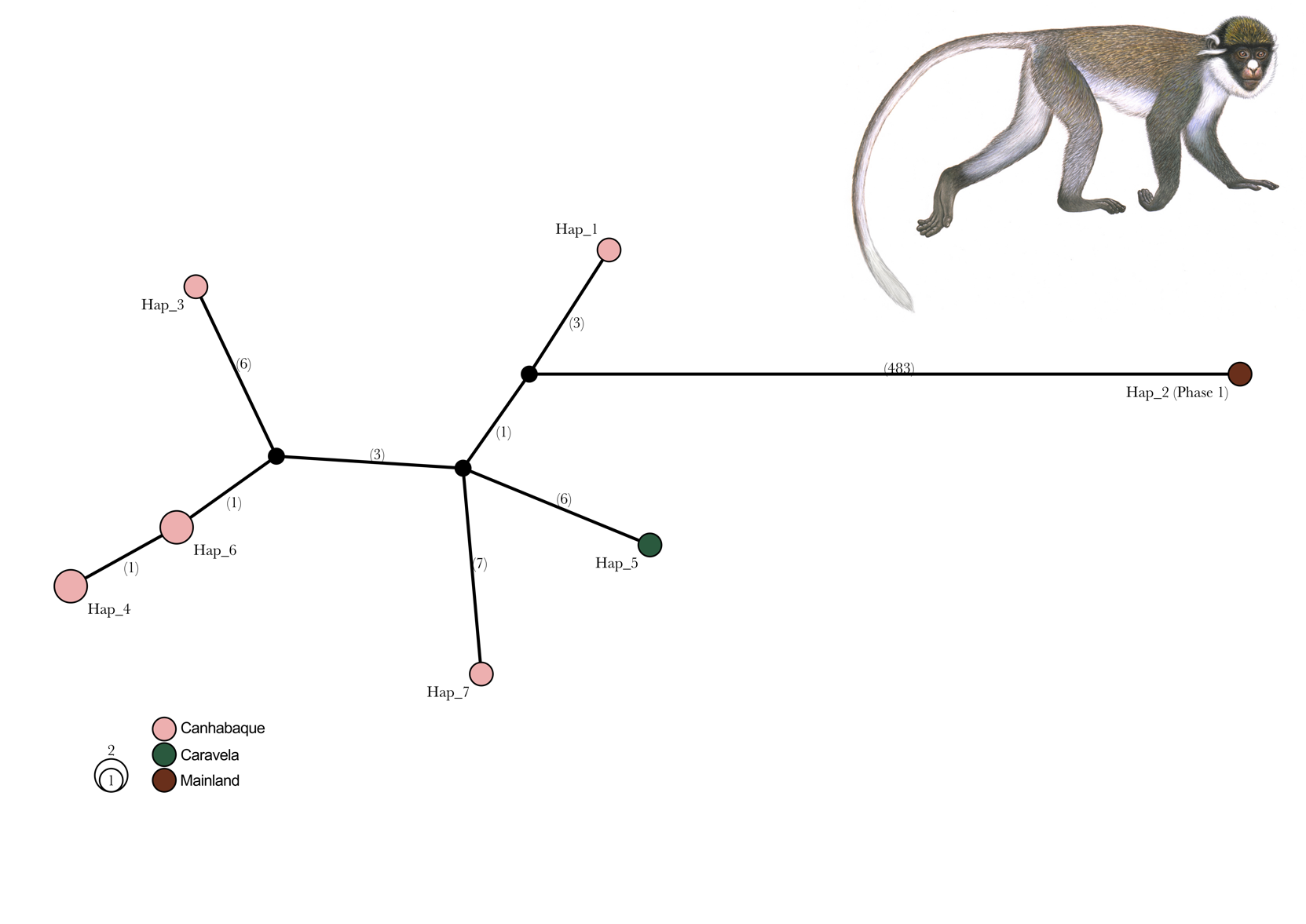

Fig. S1. Spot-nosed monkey (*Cercopithecus petaurista*) haplotype network based on whole mitochondrial sequences. Haplotype size is proportional to the number of samples and are coloured according to their sampling location in Guinea-Bissau. There are no shared haplotypes between sampling locations. Note that the spot-nosed monkey Phase_1 mainland sample is of unknow origin (Kuderna et al., 2023) but likely belongs to the Eastern spot-nosed monkey subspecies (*Cercopithecus petaurista petaurista*). The number of mutational steps between haplotypes is annotated on the branches. Illustrations copyright 2022 Stephen D. Nash / IUCN SSC Primate Specialist Group. Used with permission.

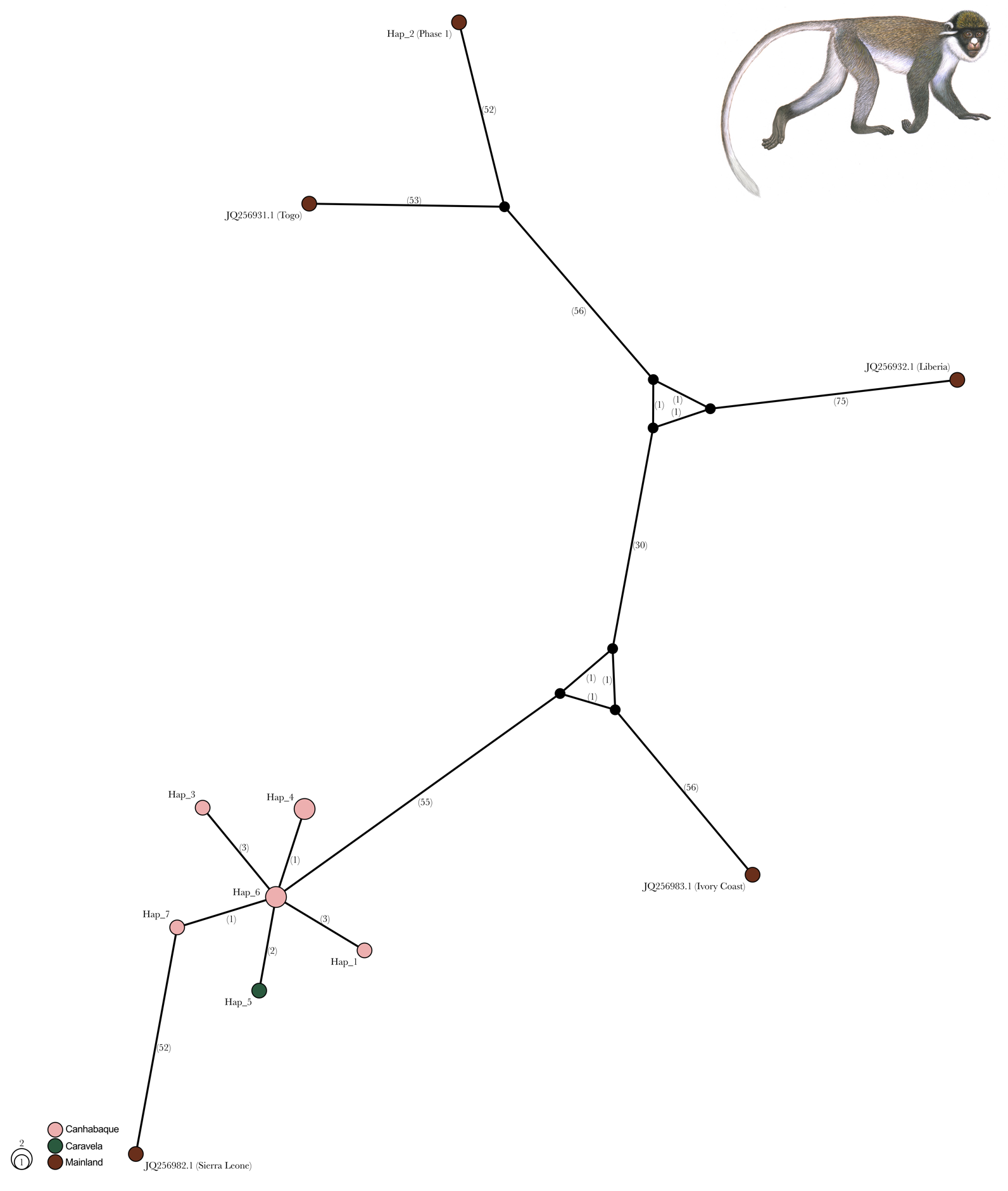

Fig. S2. Spot-nosed monkey (*Cercopithecus petaurista*) haplotype network based on whole mitochondrial sequences. Mitochondrial genomes publicly available in GenBank (https://www.ncbi.nlm.nih.gov/genbank/) were included and sampling locations were extracted from Guschanski et al., 2013. Haplotype size is proportional to the number of samples and are coloured according to their sampling location in Guinea-Bissau. There are no shared haplotypes between sampling locations. Note that the spot-nosed monkey Phase_1 mainland sample is of unknow origin (Kuderna et al., 2023) but likely belongs to the Eastern spot-nosed monkey subspecies (*Cercopithecus petaurista petaurista*). The number of mutational steps between haplotypes is annotated on the branches. Illustrations copyright 2022 Stephen D. Nash / IUCN SSC Primate Specialist Group. Used with permission.

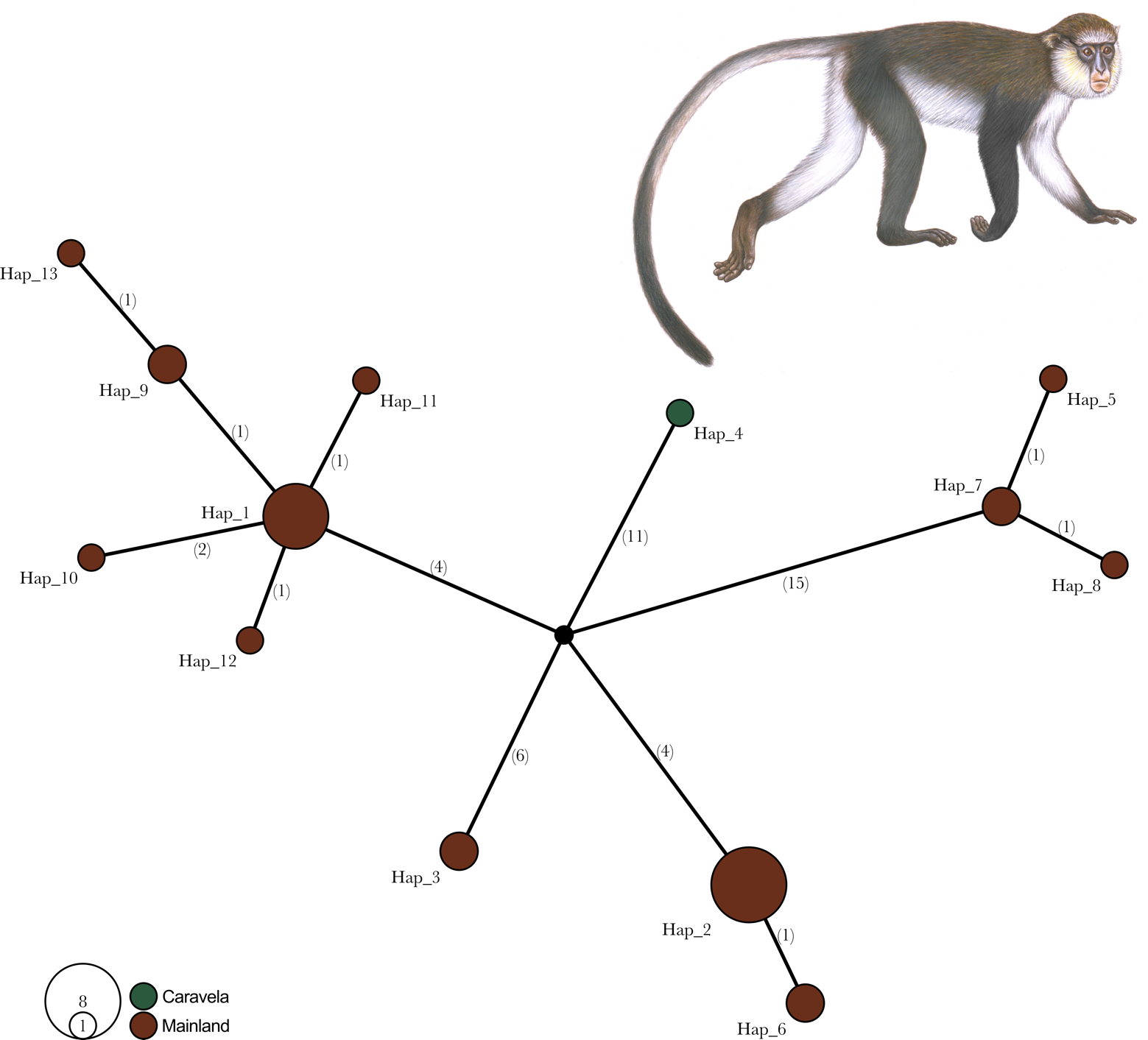

Fig. S3. Campbell’s monkey (*Cercopithecus campbelli*) haplotype network based on whole mitochondrial sequences. Haplotype size is proportional to the number of samples and are coloured according to their sampling location in Guinea-Bissau. There are no shared haplotypes between sampling locations. The number of mutational steps between haplotypes is annotated on the branches. Illustrations copyright 2022 Stephen D. Nash / IUCN SSC Primate Specialist Group. Used with permission.

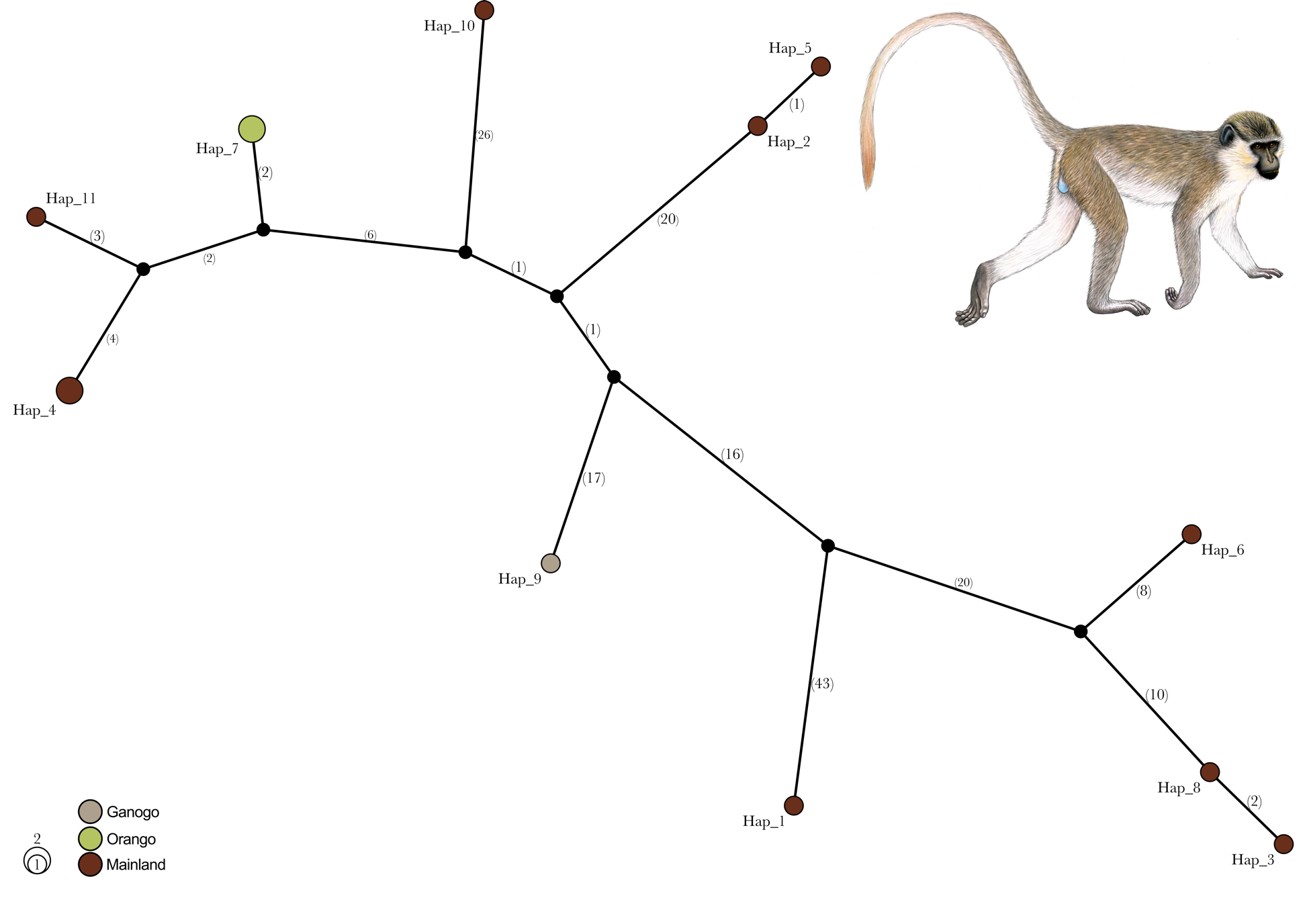

Figure S4. Green monkey (*Chlorocebus sabaeus*) haplotype network based on whole mitochondrial sequences. Haplotype size is proportional to the number of samples and are coloured according to their sampling location in Guinea-Bissau. There are no shared haplotypes between sampling locations. The number of mutational steps between haplotypes is annotated on the branches. Illustrations copyright 2022 Stephen D. Nash / IUCN SSC Primate Specialist Group. Used with permission.

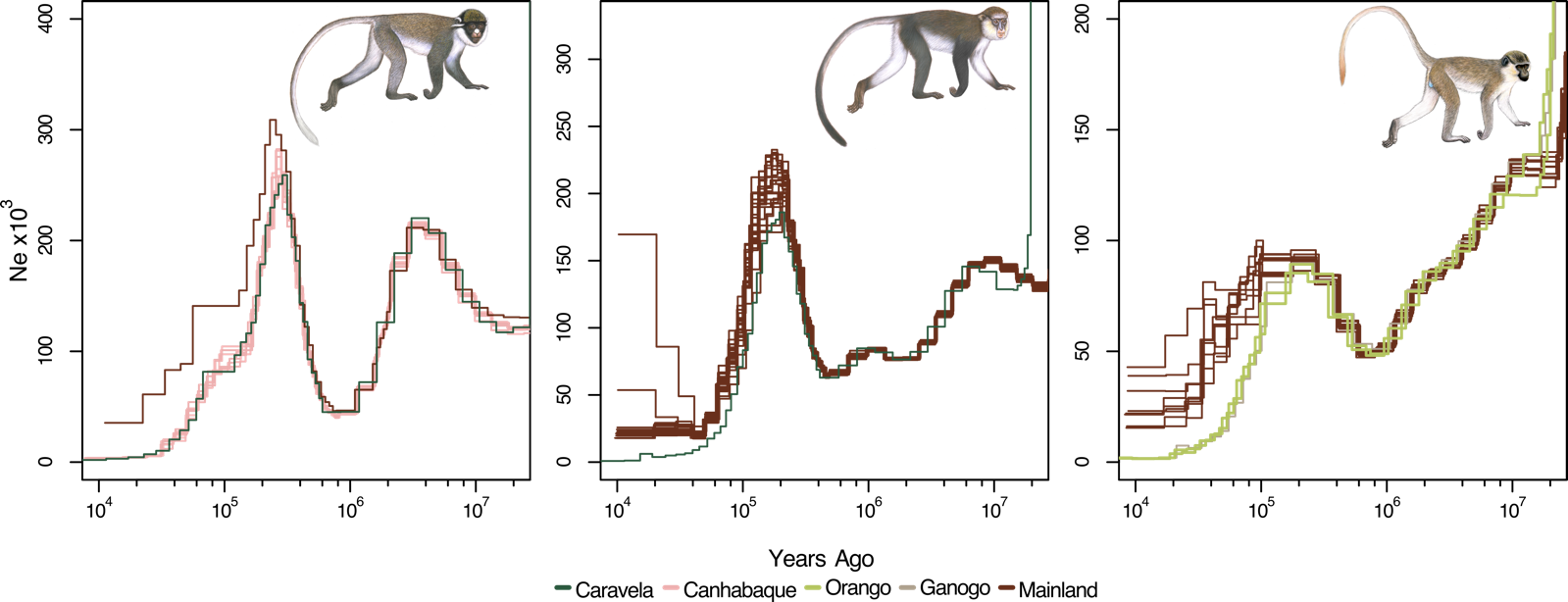

Fig. S5. Beta-PSMC analysis for all the spot-nosed monkey (*Cercopithecus petaurista*; left), Campbell’s monkey (*Cercopithecus campbelli*; middle), and green monkey (*Chlorocebus sabaeus*; right) genomes. Each line represents a distinct genome and are coloured according to the sampling location. Illustrations copyright 2022 Stephen D. Nash / IUCN SSC Primate Specialist Group. Used with permission.

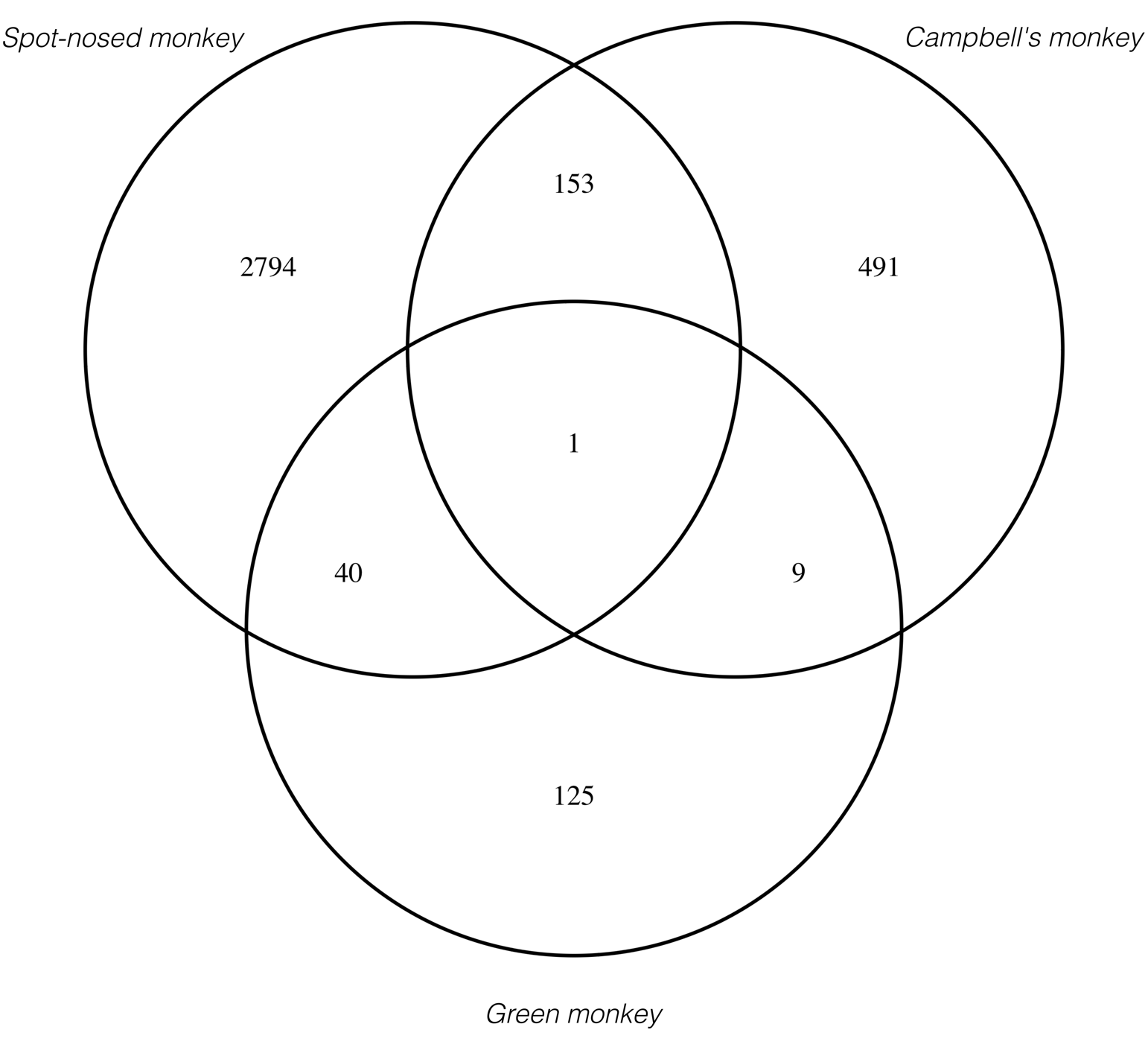

Fig. S6. Venn diagram of illustrating the number of protein-coding genes containing differentially fixed variants overlapping in the spot-nosed monkey (*Cercopithecus petaurista*: top left), the Campbell’s monkey (*Cercopithecus campbelli*; top right) and the green monkey (*Chlorocebus sabaeus*; bottom). Each line represents a distinct genome and are coloured according to the sampling location. Illustrations copyright 2022 Stephen D. Nash / IUCN SSC Primate Specialist Group. Used with permission.

| Table S1. Number of segregating sites per category of mutational effects on protein-coding genes for the three guenon species. | | | | |
| --- | --- | --- | --- | --- |
| **Spot-nosed monkey** | | | | |
|  | Caravela | Canhabaque ^†^ | Mainland | Total^‡^ |
| Synonymous | 66,289 | 68,926±803 | 71,565 | 10,7964 |
| Tolerated missense (Ex) | 29,216 | 30,181±347 | 31,165 | 50,921 |
| Deleterious missense (Ex) | 10,959 | 11,409±144 | 11,823 | 20,303 |
| LoF | 1,245 | 1,247±11 | 1,273 | 2,184 |
| **Campbell’s monkey** | | | | |
|  | Caravela | | Mainland**^†^** | Total |
| Synonymous | 66,531 | | 71,783±658 | 12,6305 |
| Tolerated missense (Ex) | 28,164 | | 30,444±272 | 62,452 |
| Deleterious missense (Ex) | 10,857 | | 11,846±130 | 25,905 |
| LoF | 1,195 | | 1,255±17 | 2,635 |
| **Green monkey** | | | | |
|  | OIG | | Mainland^†^ | Total |
| Synonymous | 69,048±1807 | | 71,426±1421 | 11,0026 |
| Tolerated missense (Ex) | 30,027±757 | | 31,341±602 | 54,681 |
| Deleterious missense (Ex) | 11,728±268 | | 12,142±231 | 22,206 |
| LoF | 1,250±29 | | 1,292±23 | 2,305 |
| † for sampling locations with N > 3, values represent the mean ± SD  ‡ total number of derived variants within species  EX missense mutations were classified as potentially deleterious or tolerated based on the experimental exchangeability score (Yampolsky et al., 2005). | | | | |
